# Efficient Multiplexing of Pollinator Metabarcodes Using Oxford Nanopore MinION Sequencing: Insights for Meadow Management from Floral Environmental DNA

**DOI:** 10.1101/2023.12.15.571580

**Authors:** Stephen E. Harris, Amelia Whitehurst, Madeline Buehrer, Sydney Lonker, Budd Veverka, Chris Nagy

**Affiliations:** School of Natural and Social Science, SUNY Purchase College, Purchase, NY, 10577 USA; Mianus River Gorge, Inc., Bedford, NY 10506, USA

**Keywords:** environmental DNA, metabarcoding, nanopore MinION multiplex sequencing, pollinator diversity, meadow management

## Abstract

Many pollinating species are experiencing declines globally, making effective, fast, and portable pollinator monitoring methods more important than ever before. Pollinators can leave DNA on the flowers they visit, and metabarcoding of these environmental DNA (eDNA) traces provides an opportunity to detect the presence of flower visitors. This study introduces an efficient, cost-effective workflow for utilizing DNA barcoding to monitor biodiversity through environmental DNA (eDNA) left on flowers from pollinators, employing the portable MinION and Nanopore sequencing. The developed method employs multiplexing with dual molecular tags on universal cytochrome oxidase 1 (COI) barcode primers. We used this approach to compare the arthropod diversity present in three meadows with different vegetation at three time points during the flowering season. The utility of eDNA metabarcoding in investigating pollinator biodiversity and interactions is highlighted through our results, revealing the presence and interactions of 65 species from 30 families. We multiplexed individual eDNA samples from hundreds of flowers and found plant-pollinator dynamics that showed differences in species richness between sampling times and meadow diversity. Comparative analyses with conventional methods showed eDNA metabarcoding’s ability to identify diverse species and ecological interactions compared to field sampling. While some DNA likely came from eggs or microscopic insects difficult to remove from flowers, traces of eDNA from various arthropods on multiple plant species confirmed the method’s applicability, promising robust ecological monitoring and research potential in the wake of global pollinator declines. This is the first reported use of MinION based nanopore sequencing to detect arthropod species from eDNA samples collected from flowers using the described affordable multiplexing method.

## Introduction

Understanding the ecological dynamics of arthropod biodiversity and their associations with the plants they feed on or pollinate is pivotal in the field of molecular ecology and biodiversity genomics, offering insights into interspecies interactions and biodiversity conservation. Pollinators, being integral components of ecosystems, are indispensable for the reproduction of most flowering plants and the production of fruits and seeds (Ollerton *et al.* 2011), thereby maintaining ecological balance and serving as proxy indicators of ecosystem health (Barsoum *et al.* 2019).

In the suburban northeast, early successional habitats such as meadows and old fields are becoming particularly rare due to forest succession, fire suppression, and human development. Many managed nature preserves therefore are now managing some land as meadows, where mowing regimes maintain an area in an early successional stage of grass or forb meadow. These meadows hopefully provide habitat for important pollinators and other native meadow species. However, maintaining a particular area in a grassland state via mowing differs substantially from the “natural” processes that kept meadows extant on the landscape in the past (Shochat *et al.* 2005). Historically, fires – either naturally-occurring or purposefully set by Native Americans, beavers (McMaster & McMaster 2001; Rosell *et al.* 2005), and other disturbances would create new meadows, while old ones would gradually succeed into forest; thus creating a dynamic “shifting mosaic” of meadows, shrublands, young forest, and mature forest across the landscape (Litvaitis *et al.* 1999). It would be helpful for land managers to know whether permanent meadows, managed by mowing, provide similar habitat value as the natural, fire-based, temporary meadows within which many species of concern evolved. Thus, techniques to efficiently and accurately sample the species assemblage of pollinators in managed meadows are needed.

The exploration of pollinator species richness and its interaction with diverse ecosystems, including meadows, is instrumental in understanding ecological equilibrium and developing conservation strategies aimed at preserving biodiversity in changing environmental landscapes (Potts *et al.* 2016). Studying the state of arthropod, and specifically pollinator richness, is especially relevant as insect diversity is declining at an alarming rate globally (Baillie *et al.* 2012; Dirzo *et al.* 2014; Rafferty 2017; Sánchez-Bayo & Wyckhuys 2019; Wagner *et al.* 2021), potentially disrupting stability across many trophic levels and ecosystems (Hallmann *et al.* 2017). In addition, the relationship between plants and their insect pollinators is the foundation of ecosystems and agriculture, with over 75% of wild plants requiring insects for pollination (Ollerton *et al.* 2011; Vanbergen & Initiative 2013). Compounding this urgency is the fact that the vast majority of plant-pollinator relationships and the extent of arthropod communities within plants is still not well understood (Thomsen & Sigsgaard 2019).

Molecular techniques, especially those leveraging environmental DNA (eDNA), have revolutionized biodiversity studies by providing a non-invasive means to study species diversity and ecological interactions, serving as an alternative to labor-intensive, potentially disruptive traditional ecological surveying techniques (Ruppert *et al.* 2019; Thomsen & Sigsgaard 2019; Baksay *et al.* 2022). eDNA allows researchers to detect species presence and abundance without direct observation, which is particularly significant in studying elusive or trace evidence of invertebrate pollinators where morphological identification is difficult (Taberlet *et al.* 2012; Thomsen & Sigsgaard 2019). eDNA metabarcoding, which entails the use of universal PCR primers to amplify species-specific fragments from mixed DNA samples, has proven instrumental in deciphering ecological communities by allowing identification of a multitude of species within environmental samples (Thomsen & Willerslev 2015).

Researchers have been using eDNA metabarcoding to study pollinator-plant interactions including identification of plant species from pollen found on the bodies of insects (Baksay *et al.* 2022), as well as the reverse, identifying visiting animal DNA from flowers (Thomsen & Sigsgaard 2019; Johnson *et al.* 2023; Newton *et al.* 2023). eDNA metabarcoding requires high-throughput sequencing, and given the unknown number of taxonomic samples, can be cost-prohibitive. In this study, we aim to take the proven concept of eDNA sequencing from flowers and apply it to affordable and portable sequencing methods. We combine eDNA metabarcoding with Oxford Nanopore Technology (ONT) sequencing, utilizing the MinION device and Flongle flow cell, to delve into pollinator species richness in disparate meadows. We will refer to this technology as Nanopore sequencing, and it enables real-time sequencing of DNA fragments, providing high-coverage and potentially long-read output for successful species identification (Jain *et al.* 2016). The nucleotide sequence of the cytochrome oxidase I (COI) gene is often unique to species and is easily amplified across animal species (Hebert *et al.* 2003). DNA barcoding is now readily being used with Nanopore sequencing for species identification (Vasiljevic *et al.* 2021). Scientists realize ONTs potential and are continually developing protocols that use miniaturized and portable equipment, and affordable home-brew methods to reduce costs to a little as 10 cents per barcode sequence (Watsa *et al.* 2020; Srivathsan *et al.* 2021; Pomerantz *et al.* 2022).

We utilize the progress made in Nanopore DNA barcoding and have developed a protocol to introduce affordable multiplexing using uniquely tagged primers (Srivathsan *et al.* 2021) and sequence eDNA samples, facilitating the simultaneous sequencing of multiple eDNA samples on one Flongle flow cell, greatly reducing the cost of sequencing. Tagging of each sample allows the segregation of sequence reads from individual flowers and enables a detailed analysis of species associations with flowers. Multiplexed eDNA Nanopore metabarcoding is a pioneering approach in investigating pollinator DNA from flowers, providing high resolution in studying ecological interactions through optimizing the throughput and sequencing needed to recover species identification from single genes (Zhou *et al.* 2013).

The exploration of insect pollinator-plant within ecosystems is crucial for understanding dynamics of biodiversity and informing natural resource management. Historically this was done using traditional ecological sampling, which often involves time-consuming and potentially disruptive techniques. Initiatives such as New York State Pollinator Protection Act and the Empire State Native Pollinator Survey are currently working to to raise public awareness while also working to understand what native insect pollinators currently inhabit New York State, and Mianus River Gorge (MRG) in Westchester County, NY is one site where the native insect pollinator biodiversity of meadows has been annually assessed since 2019 using the more traditional surveying methods. In this study, we compare an eDNA generated overall species list to the species list recorded by MRG staff and technicians in 2019 and 2021 using traditional sampling techniques. We collected wildflowers from three different meadow sites at MRG, extracted DN, and sequence amplified barcode genes to generate the eDNA list. We first show that multiplexed eDNA metabarcoding with nanopore sequencing is feasible and affordable, and second, compare the species richness results to traditional sampling methods. The combination of eDNA metabarcoding, unique molecular tagging, and Nanopore sequencing used in this study represents a feasible approach that can help advance understanding of pollinator biodiversity and meadow management. This research not only represents one of the first endeavors to employ Nanopore eDNA sequencing combined with unique molecular tagging methodologies but also highlights the immense potential of these techniques in understanding pollinator species richness in meadows.

## Materials and Methods

### Study sites and sampling

Samples for this study were collected from three distinct, managed meadow areas within the MRG Preserve, a 970 acre nature preserve located in suburban Westchester county NY, USA. (Figure 1). The meadows were all considered high flowering locations and included a meadow that had grown over an old apple orchard at Lockwoods North (LN), and two managed forb/grass meadows named Captain Woods (CW) and at Big Meadow (BM). Trapping and molecular sampling both occurred across two transects at each site. Along each transect, we defined a 1 x 1 m plot every 10 m. Traditional sampling methods (pan trapping and hand netting) were used to collect specimens using the Empire State Native Pollinator Survey (ESNPS) protocols (White *et al.* 2022) across the months of June, July, and August in 2019 and 2020 (Table 1). Insects were placed in plastic containers to be euthanized and later identified to species. Flower samples were collected for eDNA sequencing on three occasions, June 28, August 10, and September 7, 2021 (Table 3 and Supplemental Table S1 for full list of collected flowers). For eDNA sequencing, within each plot, we collected at least five flower heads from each encountered flowering plant free of visible signs of insects. Sterilized shears and nitrile gloves were used to limit cross contamination. The flower-heads were collected, identified using staff knowledge or Seek (https://www.inaturalist.org/pages/seek_app), and individually stored in sterile plastic tubes. Samples were kept in a portable cooler following collection until transportation to the lab where they were stored at -20°C.

**Fig. 1.**
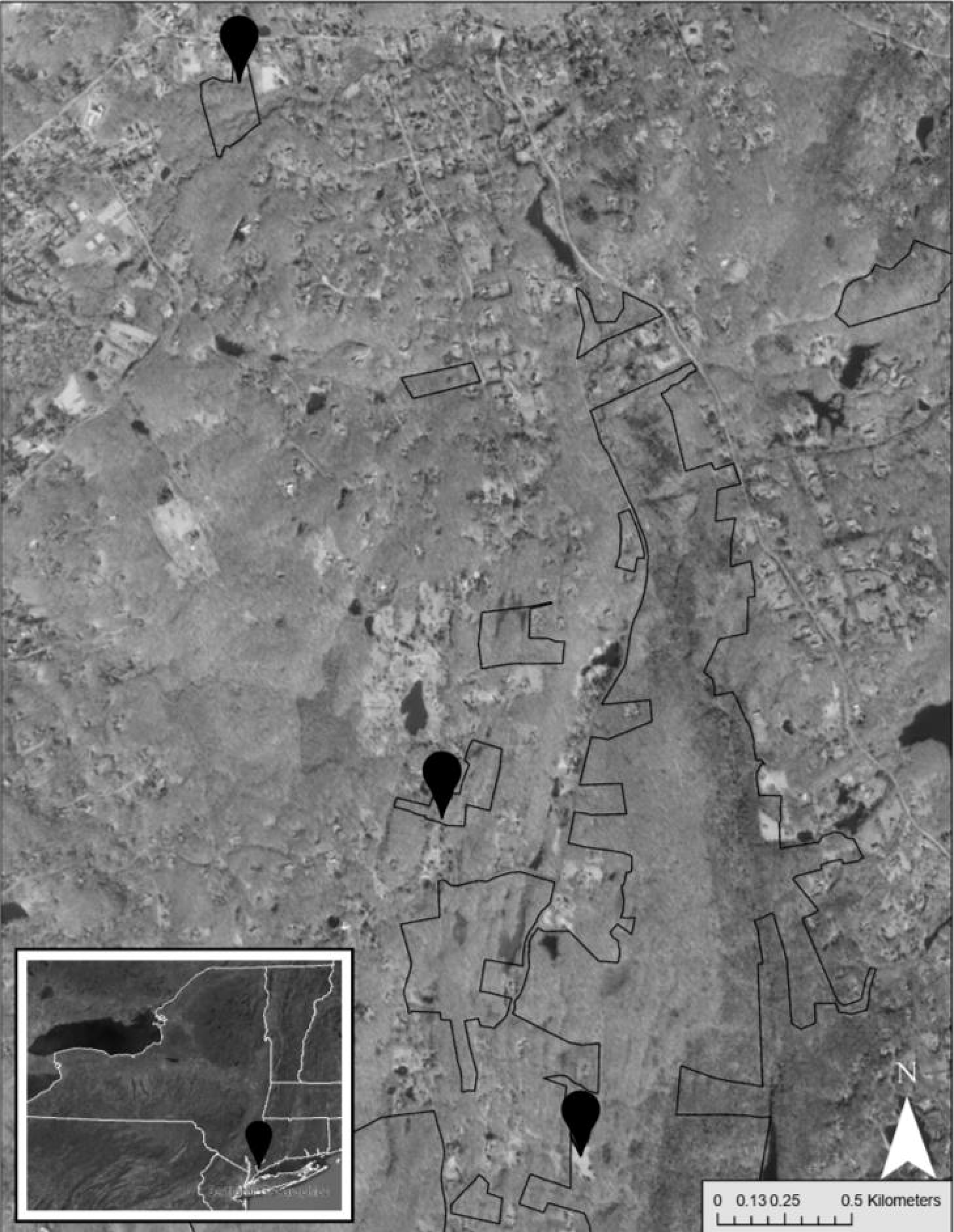
Meadows surveyed for pollinators (gray markers) in the MRG Preserve (black boundaries) in Westchester County, New York State (inset), 2022.

**Table 1:** List of taxa identified from traditional sampling methods at Mianus River Gorge. Taxa identified from hand netting and pan trapping in 2019 and 2020. Bold = also found in eDNA metabarcoding. * = focal taxa identified from ESNPS.

| Order | Family | Genus | Species |
| --- | --- | --- | --- |
| Coleoptera | Cerambycidae | *Typocerus* | *Typocerus acuticauda* |
| Coleoptera | Cerambycidae | *Typocerus* | *Typocerus velutinus* |
| Coleoptera | Buprestidae | -- | -- |
| Diptera | Asilidae | -- | -- |
| Diptera | Asilidae | *Efferia* | *Efferia albibaris* |
| **\*Diptera** | **Syrphidae** | ***Sphegina*** | ***Sphegina flavimana*** |
| *Hymenoptera | Andrenidae | *Andrena* | *Andrena nasonii* |
| Hymenoptera | Apidae | *Anthophora* | *Anthophora terminalis* |
| Hymenoptera | Apidae | *Apis* | *Apis mellifera* |
| *Hymenoptera | Apidae | *Bombus* | *Bombus gregionis* |
| *Hymenoptera | Apidae | *Bombus* | *Bombus griseocolli* |
| *Hymenoptera | Apidae | *Bombus* | *Bombus impatiens* |
| *Hymenoptera | Apidae | *Bombus* | *Bombus perplexus* |
| Hymenoptera | Apidae | *Ceratina* | *Ceratina calcarata* |
| Hymenoptera | Apidae | *Ceratina* | *Ceratina dupla* |
| Hymenoptera | Apidae | *Xylocopa* | *Xylocopa virginica* |
| Hymenoptera | Crabronidae | *Cerceris* | *Cerceris fumipennis* |
| Hymenoptera | Halictidae | *Lasioglossum* | -- |
| Hymenoptera | Halictidae | *Agapostemon* | *Agapostemon virescens* |
| Hymenoptera | Halictidae | *Agapostemon* | *Agapostemon virescens* |
| Hymenoptera | Halictidae | *Augochlora* | *Augochlora aurata* |
| Hymenoptera | Halictidae | *Augochlora* | *Augochlora persimilis* |
| Hymenoptera | Halictidae | *Augochlora* | *Augochlora pura* |
| Hymenoptera | Halictidae | *Halictus* | *Halictus confusus* |
| Hymenoptera | Halictidae | *Halictus* | *Halictus ligatus* |
| Hymenoptera | Halictidae | *Halictus* | *Halictus rubicundus* |
| Hymenoptera | Halictidae | *Lasioglossum* | *Lasioglossum coeruleum* |
| Hymenoptera | Scoliidae | *Scolia* | *Scolia dubia* |
| Hymenoptera | Vespidae | *Dolichovespula* | *Dolichovespula arenaria* |
| Hymenoptera | Vespidae | *Polistes* | *Polistes fuscatus* |
| Hymenoptera | Vespidae | *Polistes* | *Polistes peckius* |
| Hymenoptera | Vespidae | *Vespula* | *Vespula germanica* |
| **Lepidoptera** | **Hesperiidae** | ***Epargyreus*** | ***Epargyreus clarus*** |
| **Lepidoptera** | **Hesperiidae** | ***Thymelicus*** | ***Thymelicus sylvestris*** |
| **Lepidoptera** | **Nymphalidae** | ***Phyciodes*** | ***Phyciodes cocyta*** |
| **Lepidoptera** | **Nymphalidae** | ***Phyciodes*** | ***Phyciodes tharos*** |

### DNA extraction/ eDNA recovery

Five small flower heads per transect per site, or one umbel per transect per site from each flower species was/were separated into 15mL or 50mL tubes for eDNA recovery. DNA extractions were done using the DNeasy Blood & Tissue Kit (Qiagen) with modified extraction protocols described below. Each 15mL or 50mL tube received 900uL or 1800uL ATL Buffer and 100uL or 200uL Proteinase K, respectively. Samples were vortexed for a minimum of one minute to ensure maximum contact with the lysing solution for at least one minute. Tubes were incubated for three hours at 56°C with removal and vortexing for 15 seconds every hour. Specimens were then removed and vortexed for 15 seconds. Depending upon the initial volume, 700uL – 1500uL lysate was retrieved and transferred to a new 15mL tube. From here the standard DNeasy Blood & Tissue protocol was followed. DNA was eluted with 100uL of AE Buffer, incubated at room temperature for one minute, then centrifuged at 8,000g for one minute. This process was repeated a second time, for a total of 200 ul. DNA was labeled and stored at -20°C until further use. DNA yield was measured using a Qubit® 4 Fluorometer (Thermo Fisher Scientific) according to manufacturers’ protocols.

### Primer design and tagging, PCR amplification

For DNA barcoding, we used the commonly sequenced COI mitochondrial gene. eDNA is often degraded, and several trials informed our choice in choosing two primer sets (Table 2) which target different portions of the COI gene and are smaller than the traditional 658 bp amplified region (Folmer *et al.* 1994). The primer sets chosen were the forward/reverse primer pair mlCOIintF and jgHCO2198 (Table 2) which selects for 313bp (Leray *et al.* 2013), and forward/reverse primer pair ZBJ-ArtF1c and ZBJ-ArtR2c (Table 2), which selects for 157bp (Zeale *et al.* 2011). In order to multiplex samples in one sequencing run, we added a 13 base pair unique sequence tag to the forward and to the reverse COI primers, as recommended from Srivathsan *et al.* (2021). This ensured dual tagging of amplicons from each flower sample. Eight tags were created for each forward primer, and seven tags were created for the reverse primers (Table 2) creating 15 primer sequences for each primer set chosen, allowing us to multiplex 56 samples simultaneously on one Flongle flowcell. Flower samples were assigned a random forward and reverse tag_primer pair combination ensuring the dual tagging of amplicons specific to each specimen.

**Table 2.**
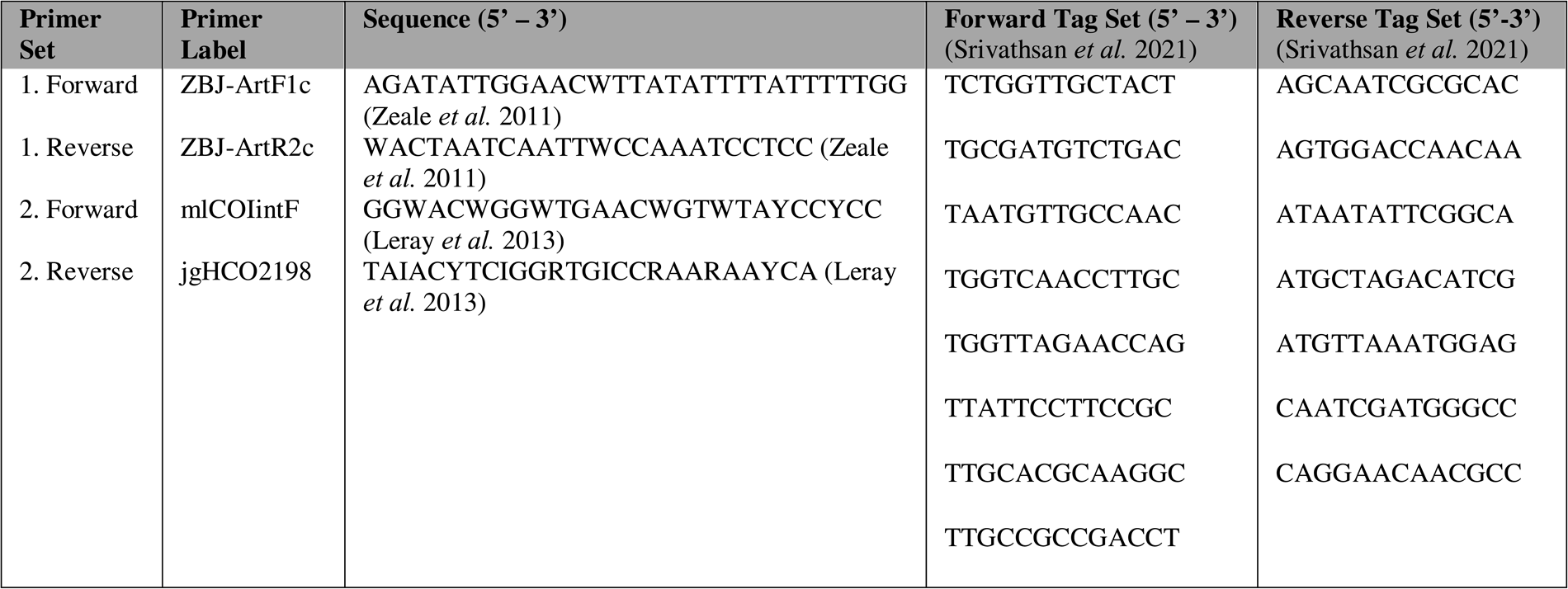
COI mtDNA primers and forward and reverse tags appended to 5’ end of primers used in this study.

### PCR amplification

The following PCR reaction was performed in 25 uL volumes. Each reaction included 12.5 uL of Amplitaq Gold™ 360 Master Mix (Invitrogen, Carlsbad, CA), 1 ul of forward primer (10 uM), 1 ul of reverse primer (10 uM), 1 ul of template DNA, 1 ul of BSA (1 mg/ml), and 5.5 of UltraPure™ DNase/RNase-Free Distilled Water (Invitrogen, Carlsbad, CA). PCR was performed twice for each specimen with uniquely tagged primer sets: mICOIintF/ jgHCO2198 and ZBJ-ArtF1c/ ZBJ-ArtR2c. Amplicons were stored at -20°C. Thermocycling parameters for primer Tag_ZBJ-ArtF1c/ Tag_ZBJ-ArtR2c1 were 95°C for 10 minutes, 55 cycles of 94°C for 30 seconds, 54°C for 30 seconds, 72°C for 60 seconds, and a final elongation of 72°C for 7 minutes. Thermocycling parameters for primer Tag_ mICOIintF /Tag_jgHCO2198 13,20 were 95°C for 5 minutes, 35 cycles of 94°C for 30 seconds, 48°C for 60 seconds, 72°C for 30 seconds, and a final elongation of 72°C for 5 minutes. Amplicons were stored at -20°C.

Sixteen randomly selected tagged post-PCR products were assessed by 2 % agarose gel electrophoresis for size verification. The post-PCR products were prepared in a 7uL solution for loading per the following: 5.0 uL amplicon DNA + 1.0 uL Blue Running Stain + 1.0uL Midori Green Advance DNA Stain (Nippon Genetics).

### Library preparation and Nanopore sequencing

We ran six sequencing runs, three for primer Tag_mICOIintF / Tag_jgHCO2198 and three for primer Tag_ZBJ-ArtF1c / Tag_ZBJ-ArtR2c for June, August, and September. DNA extracted from each flower species was amplified with its own unique forward and reverse tag for up to 56 uniquely dual-tagged amplicon pools. There were no overlapping tag combinations within each sequencing run. Sequencing libraries were prepared by pooling amplified DNA from between 25 samples (1 ul each) and 43 samples (0.6 ul each) from each dual-tagged post-PCR product for a given month and primer pair then assessed to ensure DNA concentration ≥ 100ng / 25uL using a Qubit® 4 Fluorometer (Thermo Fisher Scientific). Each pooled amplicon library was prepared with the Oxford Nanopore Technologies (ONT) Ligation Sequencing Kit (SQK-LSK109), NEBNext® Companion Module for Oxford Nanopore Technologies® Ligation Sequencing (E7180S, New England Biolabs), and ONTs Flongle Sequencing Expansion Kit (EXP-FSE001) according to the manufacturer protocols. Final sequencing libraries were loaded onto a Flongle™ Flow Cell R9.4.1 (FLO-FLG001). Sequencing was performed using the Oxford Nanopore Technologies MinION™ M1Kb device and MinKNOW (v23.04.06).

### Sequencing analysis

Sequences were basecalled using ONT Guppy v2.3.7 on the raw fast5 files using the high accuracy (HAC) model. Demultiplexing was performed with ONT Barcoder v0.1.9 (Srivathsan *et al.* 2021). The primer sequences with tag assignments was used as input. Remnant primer sequences, tags, and adapter sequences were then removed using porechop v0.2.3 (Wick *et al.* 2017). Quality control of sequences was done with NanoFilt v2.2.0 (De Coster *et al.* 2018), filtering reads with a q ≥ 10 and minimum read length between 300 bp and 800 bp for mICOIintF/ jgHCO2198 and between 150 bp and 500 bp for ZBJ-ArtF1c/ZBJ-ArtR2c. The resulting sequences were then compared against a COI database containing all COI DNA sequences ≥ 300 bp in length downloaded from the NCBI nt database on 05/01/2023. Sequences were given taxonomic assignment using BLASTn v2.9.0 (Altschul *et al.* 1990) using an e-value of 1e-15 and max target sequences of one. Taxonomic classification was assigned using R package, *taxonomizr*. BLAST results were further filtered by keeping only hits ≥ 100 bp in match length, ≥ 98% identity match, and occurring ≥ 10 times. Nanopore fastq files q ≥ 10 are are available from the Dryad Digital Repository (https:doi:xx.xxx/dryad.XXXX)

### Data analysis

Plot-by-plot capture data was not available from the 2019 and 2021 trapping surveys, so we were only able to compare the final species list from each site across the two methodologies. All species richness analyses were performed in RStudio v2023.09.0-463 (RStudioTeam 2020). Species richness and various diversity measures (Shannon and Simpson) for each site (LN, CW, BM) and sampling time (June, August, September) were calculated using the R package *vegan* specaccum from the R package *vegan* v2.4-4 (Oksanen 2017). Rarefaction curves for all nine sequencing pools were made using counts data of number of taxa with the R package *vegan* v. 2.4-4 (Oksanen 2017). We also used *vegan* to perform an ordination analysis, specifically a PCA, to see if pollinator communities were differentiated based on sampling location or sampling period. Finally, to visualize links between plants and pollinators, we used the R package *bipartite* (Dormann *et al.* 2008). Finally, we compared species by species between traditional sampling survey and our eDNA approach, and then tallied the number of taxa found overlapping with the important focal pollinator taxa from the ESNPS report.

## Results

### Sequencing and BLAST results

We sequenced a total of 1,569,917 reads ≥ q 10 from six Flongle flow cell runs. There was some variation in sequencing depth across samples 4,899 ± 574 reads (mean, SEM). After demultiplexing, setting minimum and maximum length requirements, and removing adapters, there were 695,647 reads.

Initial BLASTn results found 569 unique species (Dryad Digital Repository, https:doi:xx.xxx/dryad.XXXX). After requiring a minimum match length of 100 bp and 98 % Identity, we found 65 species from 30 families and 12 orders (Supplemental Table S2). We considered species with ≥ 10 individual BLAST hits within an individual flower as high confidence hits (Table 3).

**Table 3.**
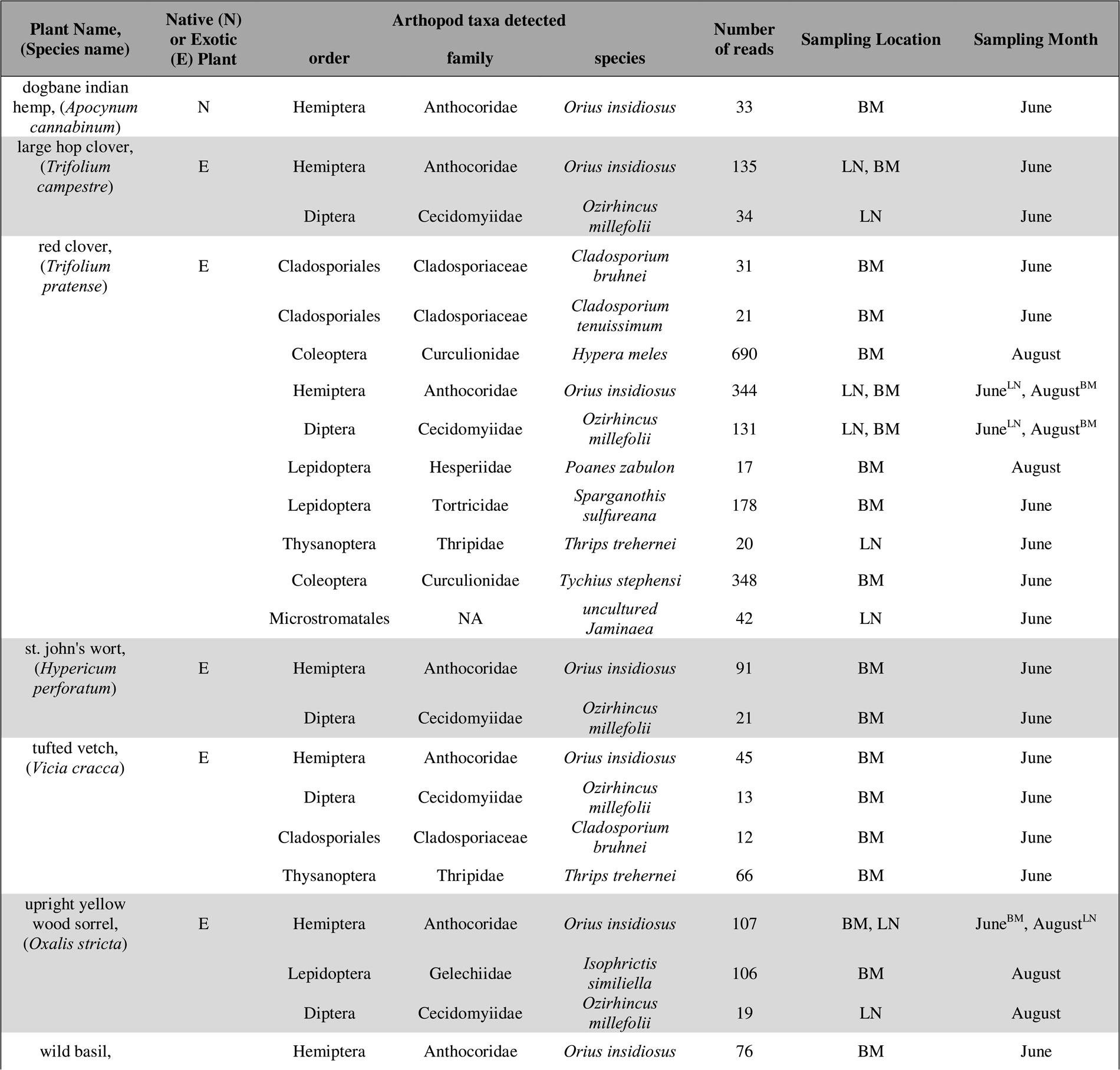
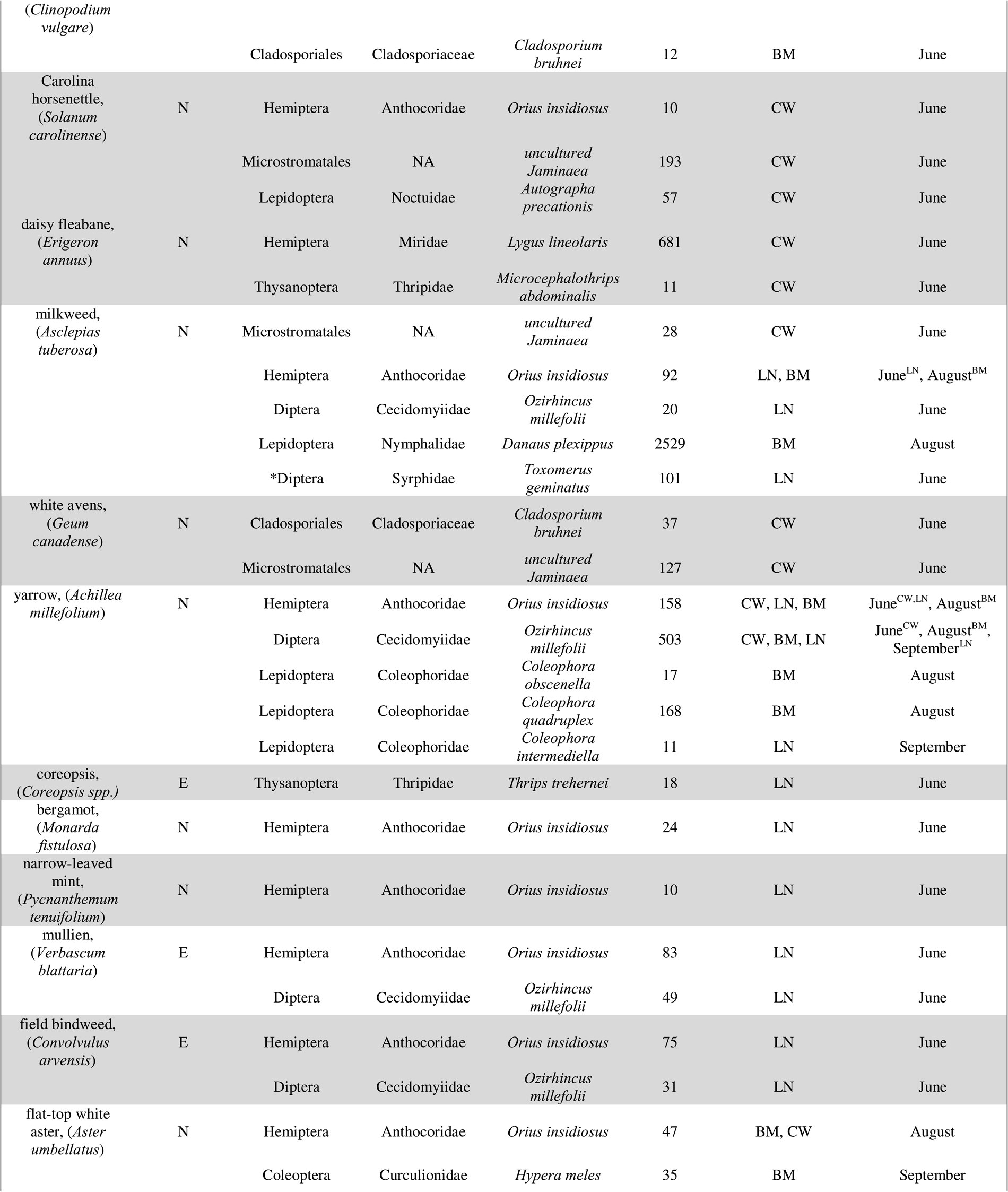
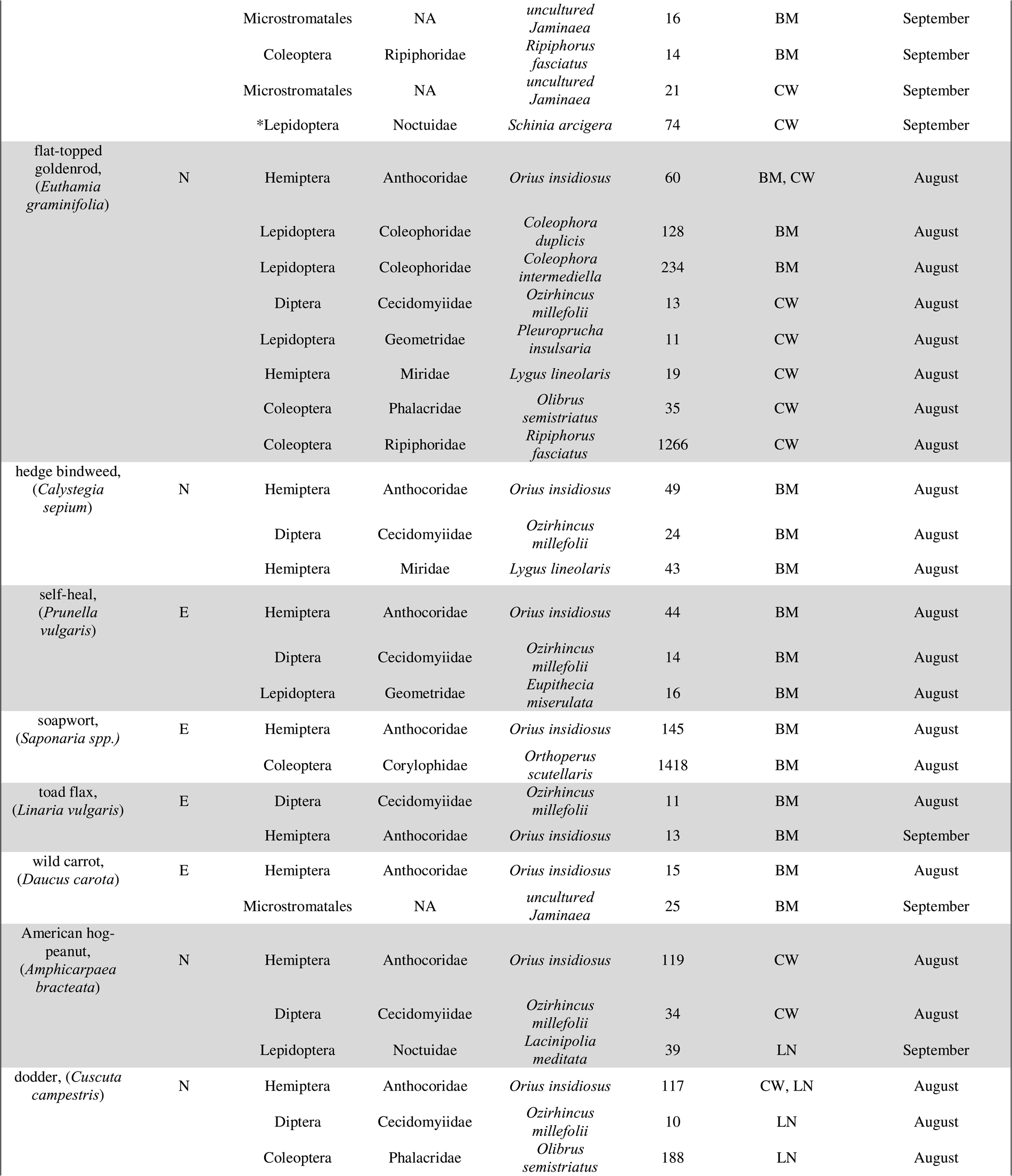
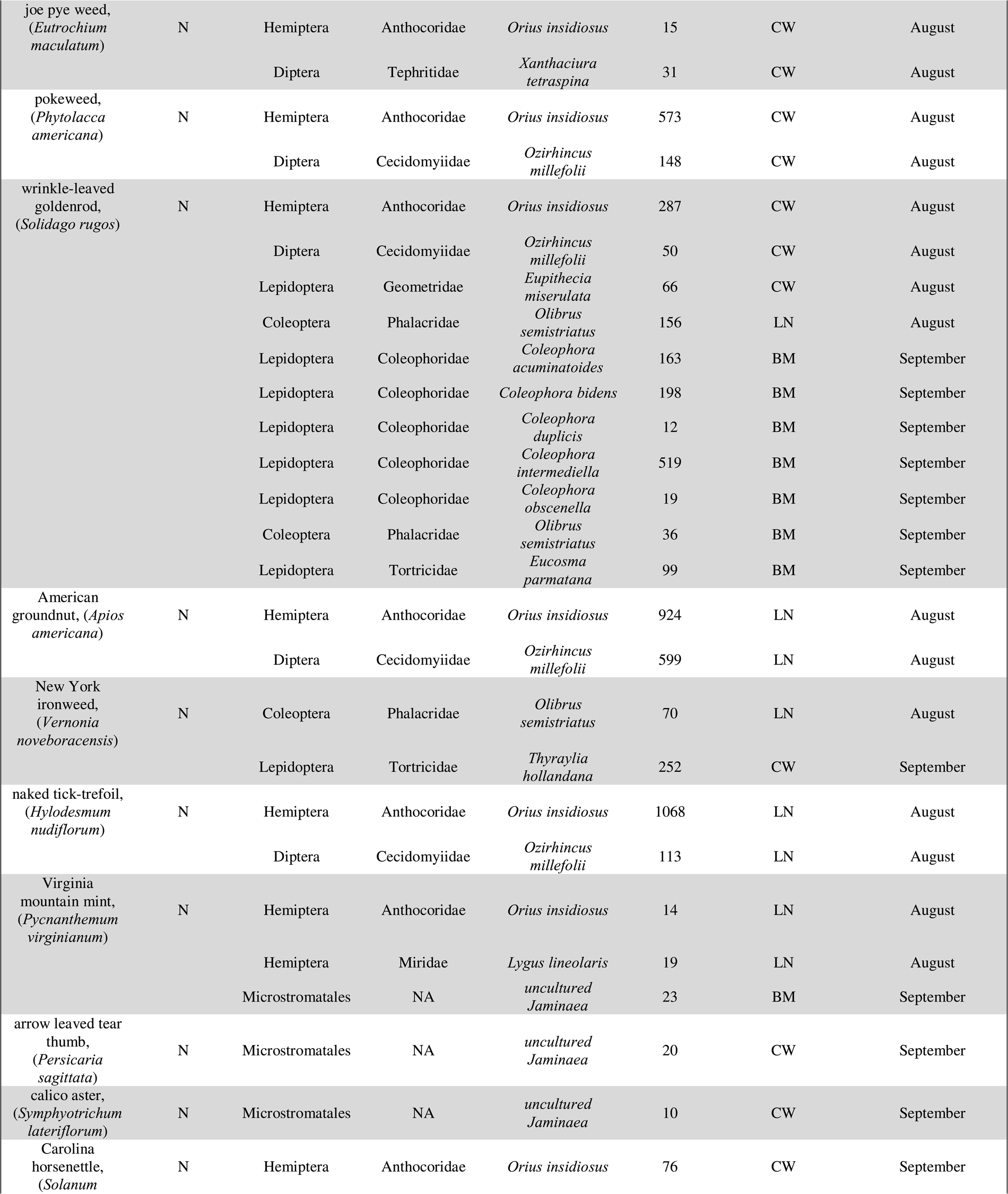
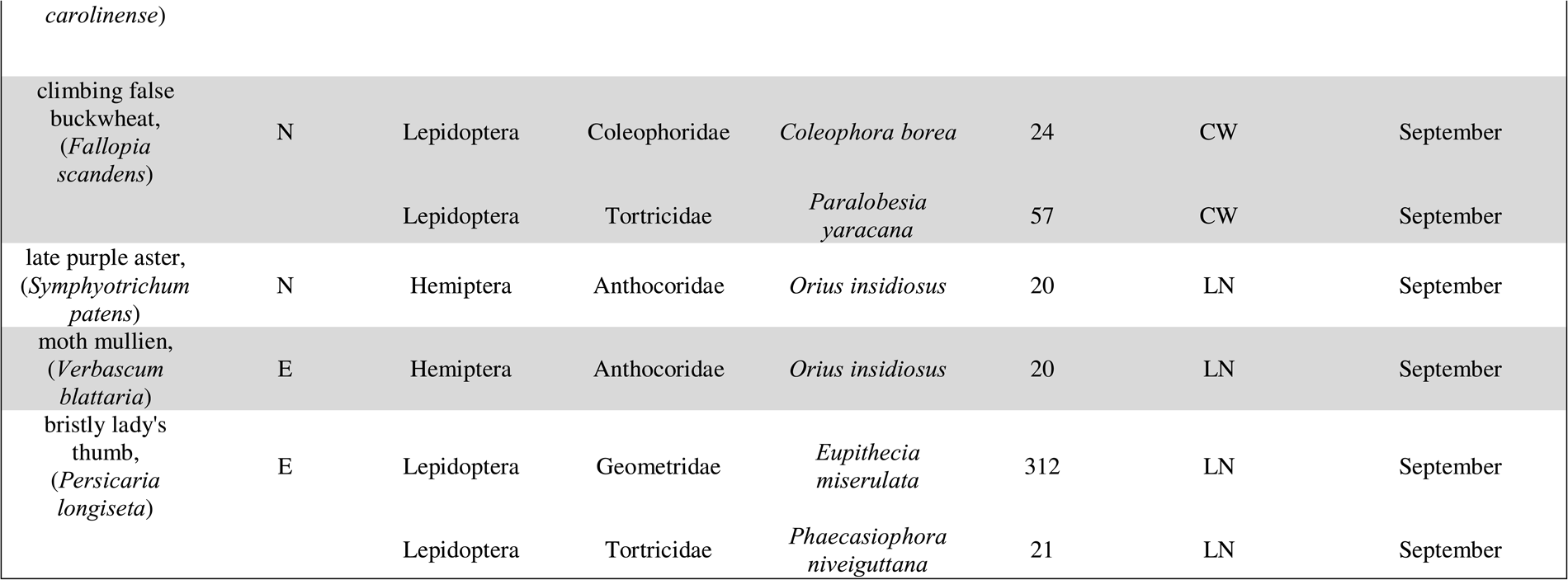
High confidence species hits from BLAST search with 98% ID, minimum length 100 bp, and occurrence ≥ 10 within an individual flower sample. BM = Big Meadow, CW = Captains Woods, LN = Lockwoods North. Native = N, Non-native/Exotic = E, Unknown Status = U. When ^LN,CW, or^ ^BM^ are added to sampling month, this specifies when and where an insect was identified. * in the column ‘order’ = focal taxa identified from ESNPS. Native versus exotic status was determined using the New York Floral Atlas from the New York Floral Association (https://newyork.plantatlas.usf.edu/).

### Plant species

We collected 58 flower species in total and successfully extracted animal DNA and identified animal species from 41 flower species (Table 3 & Supplemental Table S1). There were 14 flowers that were found at multiple sites and/or times. Yarrow (*Achillea millefolium)*, was the only plant found in multiple sites across all three sampling months (June, August, September). Flat-top white aster (*Aster umbellatus)* was also prevalent at multiple sites in August and September. New York ironweed (*Vernonia noveboracensis),* upright yellow wood sorrel *(Oxalis stricta),* milkweed *(Asclepias tuberosa),* Carolina horsenettle *(Solanum carolinense),* large hop clover *(Trifolium campestre),* toad flax *(Linaria vulgaris),* Virginia mountain mint *(Pycnanthemum virginianum),* wild carrot *(Daucus carota),* and wrinkle-leaved goldenrod *(Solidago rugos)* were also found at multiple sites and in at least two of the sampling months.

### Animal species identified from eDNA

We identified multiple species beyond arthropods in our dataset. There were several *Home sapiens* sequence reads, and we counted these as contamination. There were also reads matching to fungal species, birds, and domesticated cattle (Dryad Digital Repository, https:doi:xx.xxx/dryad.XXXX). None of these sequences occurred more than 10 times so it is difficult to be confident in their matches and they were not included as high confidence hits. Only arthropods and one fungal species met the threshold for high-confidence hits, representing 35 species (Table 3). The arthropods included species from two families of bugs, one family of midges, five families of moths, three families of beetles, one family of weevils, two families of butterflies, one family of hover flies, one family of fruit flies, and one family of thrips (Table 3). The minute pirate bug, *Orius insidiosus,* from the order Hemiptera was the most abundant species identified, occurring in 41 separate flower samples across all sites and sampling times. The gall midge, *Ozirhincus millefolii,* of order Diptera was found in 22 flowers. Members of the order Coleoptera were common, with the flower beetle, *Olibrus semistriatus,* occurring in 5 samples, the weevil, *Hypera meles,* found on two clover samples, and the wedge-shaped beetle, *Ripiphorus fasciatus,* occurring across samples as well. Lepidoptera was well represented with four moth species being found across multiple flowers, sites, and time periods (Table 3). There were also other Hemiptera and Thripidae, *Lygus lineolaris* and *Thrips trehernei*, respectively.

### Stastical analyses and plant/arthropod community comparisons

There were measurable differences in communities between sites and sampling times. We used species richness (Shannon and Simpson’s) to look at patterns of α-diversity (Table 5). BM had the highest measures of insect diversity compared to CW and LN. August had higher levels of insect species richness compared to June and September, and the general pattern was lower insect diversity in June, a peak in August, with a slight decrease in diversity in September (Table 4).

**Table 4.** Total arthropod species richness (from counts) calculated for each site, then split by native plant arthropod richness versus exotic plant arthropod richness, and then by each sampling time. Shannon’s (Typically ranges from 1.5 – 3.4, higher = more diverse) and Simpson’s (closer to 1 = greater diversity) Index of Diversity for each site.

| Location,<br>Date | Total<br>Shannon's | Native<br>Shannon's | Exotic<br>Shannon's | June<br>Shannon's | August<br>Shannon's | September<br>Shannon's | Total<br>Simpson's | Native<br>Simpson's | Exotic<br>Simpson's | June<br>Simpson's | August<br>Simpson's | September<br>Simpson's |
| --- | --- | --- | --- | --- | --- | --- | --- | --- | --- | --- | --- | --- |
| Big<br>Meadow,<br>June | 2.40 | 1.88 | 1.91 | 1.76 | 1.80 | 1.98 | 0.86 | 0.77 | 0.80 | 0.77 | 0.76 | 0.78 |
| Captain's<br>Woods,<br>June | 2.05 | 1.82 | 1.14 | 1.39 | 1.33 | 1.73 | 0.82 | 0.78 | 0.58 | 0.65 | 0.65 | 0.75 |
| Lockwood<br>North,<br>June | 1.45 | 0.79 | 0.16 | 1.44 | 0.95 | 1.46 | 0.65 | 0.35 | 0.051 | 0.65 | 0.52 | 0.67 |

The cluster analysis using PCA showed that arthropod communities identified from eDNA were largely segregated by sampling time, forming clear clusters (Figure 2). Within each time cluster, BM was farther away and with more distinct species than CW and LN; for each time point, CW and LN clustered closer together than BM (Figure 2).

**Fig. 2.**
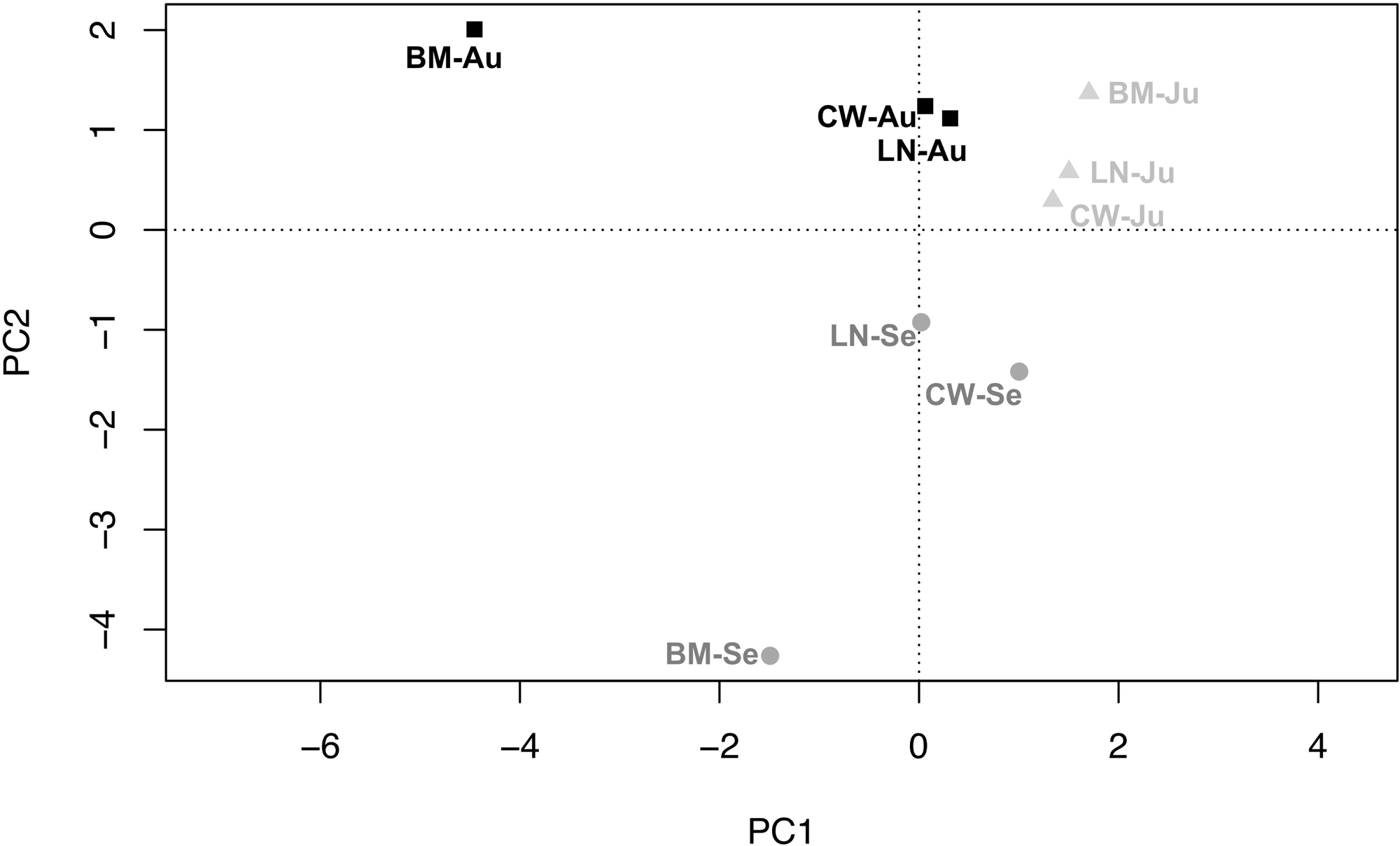
Principle Component Analysis (PCA) for COI for each site and sampling time period. BM-Au = Big Meadow – August, CW-Au = Captain’s Woods – August, LN-Au = Lockwood North – August, BM-Ju = Big Meadow – June, CW-Ju = Captain’s Woods – June, LN-Ju = Lockwood North – June, BM-Se = Big Meadow – September, CW-Se = Captain’s Woods - September, LN-Se = Lockwood North – September.

The insect orders, Lepidoptera, Diptera, Coleoptera, and Hemiptera were the most common across libraries and generated the most sequence results (Figure 3). Generally, large flowers or those with large numbers of flowers per plant contained the most insect species diversity, including goldenrod (*Solidago rugos, Euthamia graminifolia*), yarrow (*Achillea millefolium*), and milkweed (*Asclepias tuberosa*) (Table 3, Figure 3). A sample of wrinkle-leaved goldenrod (*Solidago rugos*) had 7 insect species found, the highest number of recovered species from a single flower specimen (Figure 3).

**Fig. 3.**
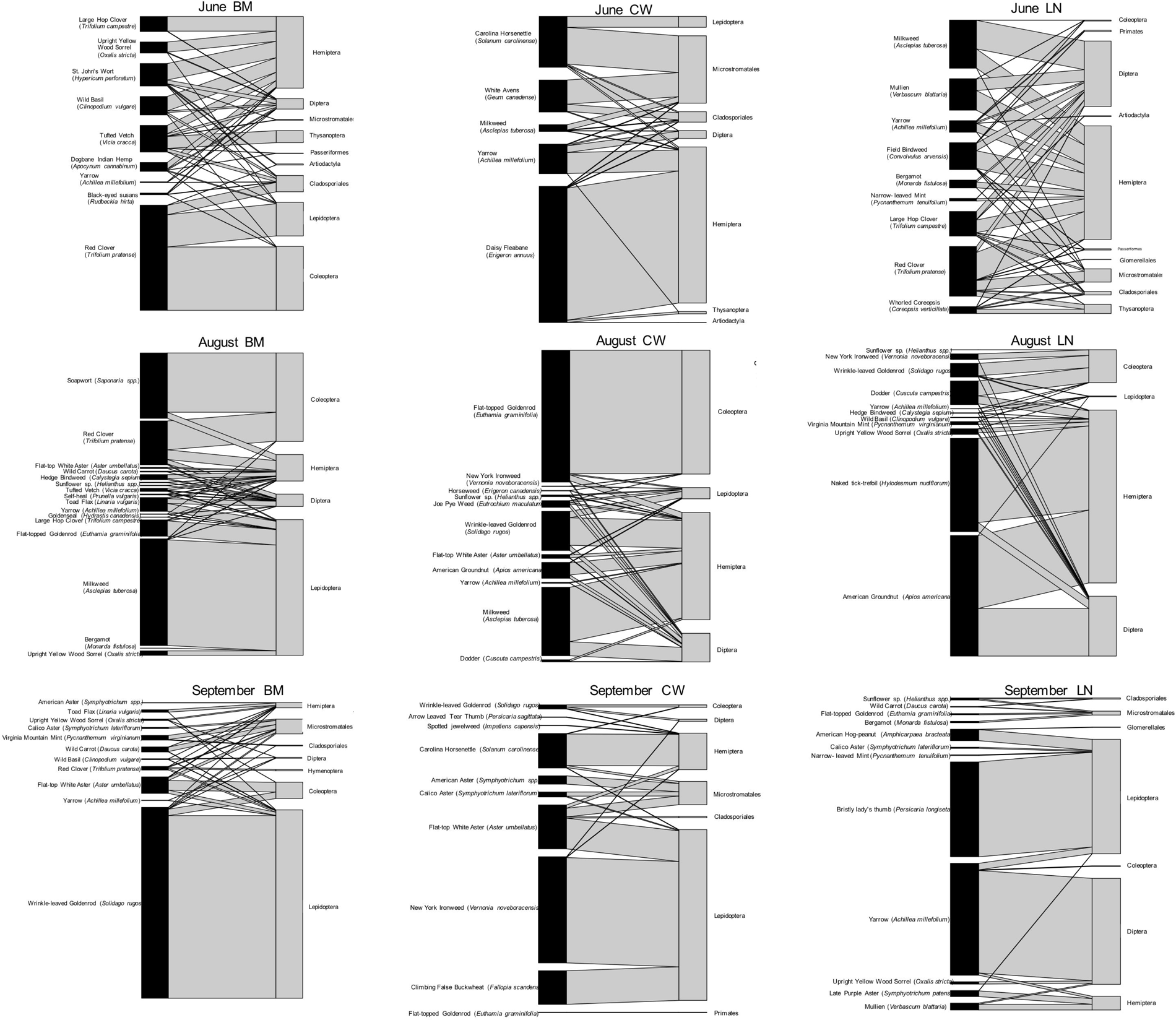
Bipartite plots for COI for all sampling sites and times showing arthropod taxa (right of each graph) found on each plant taxa (left of each graph). Plants are labelled as genus or species. Arthropods are labelled as family. The size of the connection between family and plant is proportional to the number of DNA sequences recovered for that species from a given flower. BM = Big Meadow. CW = Captain’s woods. LN = Lockwood North.

The most abundant insect species found were those which spend multiple life stages on a single host plant like plant bugs (*Orius insidious,* found in 41 flowers and *Lygus lineolaris*, found in 4 flowers), gall midges (*Ozirhincus millefolii*, found in 22 flowers), flower beetles (*Olibrus semistriatus*, found in 5 flowers), and thrips (*Thrips trehernei,* found in 3 flowers) (Table 3). BM had 53 insect species recovered from eDNA, the highest number of recovered species from plants, followed by 35 species from CW, and 37 species for LN (Table 3). BM also had the highest number of non-native plants compared to CW and LN (Table 3). A particular arthropod species was often recovered from both native and non-native plants, but there were much higher incidences of arthropods being found only in native plants and not in non-native plants. There were 22 arthropod species that were found only in native plants versus 6 arthropod species found only in non-native plants (Table 3), and 6 species found in native and exotic plants.

### Comparison between eDNA and traditional sampling

We identified seven orders, 13 families, and 36 species (after requiring > 10 reads) compared to three orders, nine families, and 33 species from traditional sampling (Table 1 and Table 3). eDNA did detect three more species than traditional sampling, and this came from more diversity in families. We did not find any overlapping species between the two methods, only species from three overlapping families, including butterflies in the family Hesperiidae, butterflies in the family Nymphalidae, and flies in the family Syrphidae. This represents 10.3% of families found in both datasets. Species that were missed by eDNA were mainly from the order Hymenoptera representing large temporary visitors to flowers like bees and wasps that may not leave much eDNA. Traditional sampling found six species from three families included in the Empire State Native Pollinator Survey’s list of important focal taxa including four species of bumble bee, *Bombus spp.,*a fly in the *Syrphidae* family, and the bumped miner bee, *Andrena nasonii.* eDNA found two arthropod species from two families included in the ESNPS, the eastern calligrapher S*yrphid* fly, *Toxomerus geminatus,* and the flower moth, *Schinia arcigera.* (Supplemental Table 2 and 3).

## Discussion

Arthropods are one of the most abundant groups on earth (Stork 2018) and the relationship they form with plants forms the foundation of ecosystem functioning and success (Ebeling *et al.* 2018). Arthropods, representing a sprawling diversity, and their interaction with plants drive important ecosystem functions like pollination, herbivory, nutrient cycling, and pest-control, and decades of research now show a direct link between arthropod and plant richness and ecosystem success (Tilman *et al.* 2014; Ebeling *et al.* 2018; Katumo *et al.* 2022). However, recent studies have shown a concerning decline in arthropod populations, with climate and land-use changes being pivotal drivers of biodiversity turnover in arthropod communities (Sánchez-Bayo & Wyckhuys 2019).

Monitoring arthropod diversity and documenting pollinator-plant interactions *in situ* are fundamental for understanding and conserving biodiversity. Our study showed that multiplexed eDNA Nanopore sequencing can supplement traditional monitoring methods that are labor-intensive, sometimes invasive, and which may miss the depth of the true species assemblage. The advent of environmental DNA (eDNA) and metabarcoding technologies has allowed an expansion in biodiversity monitoring, unveiling novel interactions between arthropods and plants (Thomsen & Sigsgaard 2019; Yoneya *et al.* 2023). Particularly, portable ONT Nanopore sequencing facilitates real-time, high-throughput sequencing of eDNA for metabarcoding studies (Egeter *et al.* 2022).

Identifying key native and present or competing non-native pollinators, identifying their respective use of – and thus the ecological value of – native versus non-native plants is vital for understanding ecological dynamics and conservation efforts. Native plants often host more specialized plant-pollinator visitation networks than non-native plants, which tend to be used solely by generalist pollinators (Seitz *et al.* 2020). However, understanding the mechanisms by which non-native plants attract pollinators and impact native-plant pollinator networks is still not well understood, especially regarding how invasion pollinators and plants can disrupt native interactions (Parra-Tabla & Arceo-Gómez 2021; Abdallah *et al.* 2021). An affordable, portable, and non-invasive approach, as presented here, is a useful additional tool to studying plant-pollinator community networks at a global scale in the face of intensifying climate change.

### Arthropod detection from eDNA

Despite lower output and coverage from ONT Nanopore sequencing compared to Illumina sequencing, we still recovered a high proportion of taxonomic diversity. Rarefaction curves showed that for several sequencing libraries, sequencing depth was sufficient, however, greater sampling effort would increase detected diversity, especially in June and September (Figure 4).

**Fig. 4.**
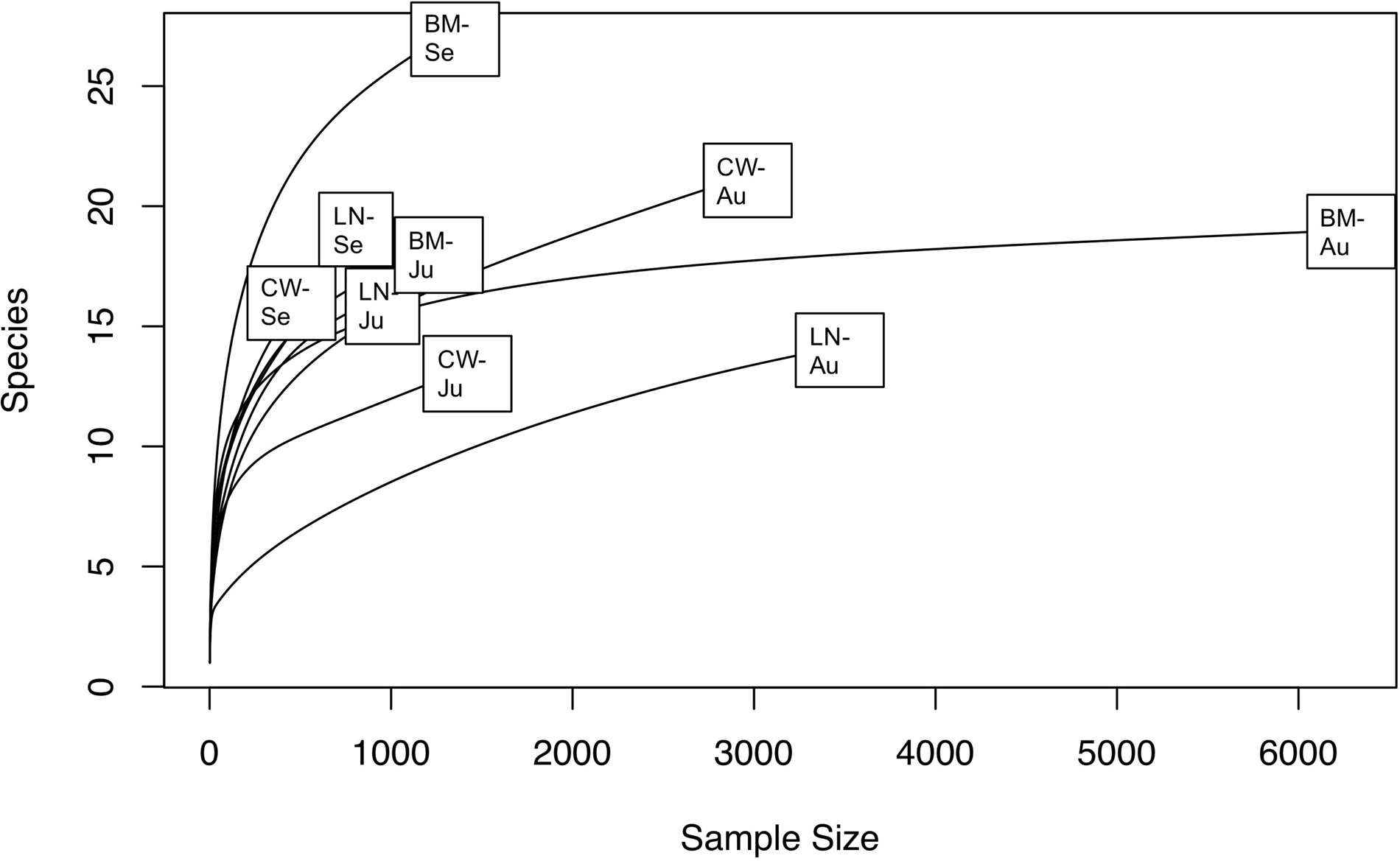
Rarefaction curves for each sequencing library. Sample size refers to the number of sequencing reads and Species is the number of species identified from eDNA. BM-Au = Big Meadow – August, CW-Au = Captain’s Woods – August, LN-Au = Lockwood North – August, BM-Ju = Big Meadow - June, CW-Ju = Captain’s Woods – June, LN-Ju = Lockwood North – June, BM-Se = Big Meadow – September, CW-Se = Captain’s Woods – September, LN-Se = Lockwood North – September.

Additionally, increased sampling of different flower species and running PCR replicates would likely also increase species recovered (Thomsen & Sigsgaard 2019). Many of the most abundant species from our analysis live on flowers in high abundance. Our results indicate that species diversity tends to vary more across season than across site. (Figure 2). The Big Meadow site is the exception, showing differentiation by both month and from the other two locations. All three sampling locations were considered high quality meadows, but BM is substantially larger than the other two and is the most ecologically intact of the three: more flower species were recovered from BM overall and in each sampling month.

Similar to Thomsen and Sigsgaard (2019) and Johnson *et al.* (2023), the majority of species identified and the bulk of the contributing reads came from small cryptic pollinators or pests, like thrips, bugs, gall midges, weevils, and larvae that feed on and inhabit flowering plants (Table 3). For example, we found the common insidious flower bug, *Orius insidiosus*, an efficient predator of thrips, on the greatest number of different flowers (44, Table 3). We also found the gall midge, *Ozirhincus millefolii,* on 22 different flowers and which may ultimately contribute to pollination (Dorchin *et al.* 2015). The thrip, *Thrips trehernei*, was found in many different samples, not surprising as it lives and breeds directly in flowers. We found similar high occurrence of the flower beetle, *Olibrus semistriatus*, one of ∼30 species that live on flowers feeding on fungus or flower heads themselves. In many case, insect species’ were identified from eDNA recovered from the flower species which serves as their primary host or food source. We found eDNA from the moth, *Coleophora intermediella*, on goldenrod, whose larvae feed on goldenrod flower seeds. We found eDNA from another moth, *Coleophora duplicis,* on aster and goldenrod flowers, whose larvae feed on asters. eDNA from the weevil, *Hypera meles,* was found on several flowers including red clover, and it feeds on red and white clovers. Additionally, we found eDNA from the moth, *Coleophora obscenella,* on common yarrow, again, whose larvae feed directly on yarrow samples. One of the more interesting finds was identifying a wedge-shaped beetle, *Ripiphorus fasciatus*;, an unusual parasitic coleopteran that parasitizes bees, including important pollinators from the families Halictidae and Apidae (Majka *et al.* 2006), two groups which were only found through traditional sampling methods (Table 1). Very little is known about the life history of the wedge-shaped beetle, or the genus *Ripiphorus* due to their complicated lifecycle; bees pick up larva from flowers and the larva mature in the hymenopteran nest. The adult life stage is only 1-2 days (Majka *et al.* 2006). Eggs are laid in the flowers and hatch as the flowers bloom, synchronizing with the arrival of host bees. DNA, likely from eggs or early larvae, was identified from flat-top white aster, flat-topped golden rod, and traces were found in toad flax, joe pye weed, hedge bindweed, and carolina horsenettle (Supplemental Table 2). This may be the first recorded observation of flower hosts for *Ripiphorus spp.* and can help in understanding the distribution of *Ripiphorus* and its impact on bees. This evidence suggests that our multiplexing approach, and its use for metabarcoding with Nanopore sequencing on the MinION, can successfully identify arthropod species interacting with flowers and can provide novel insights into especially complicated insect species phenology.

### Multiplexed eDNA nanopore metabarcoding vs. traditional sampling

We identified more arthropod species compared to traditional sampling methods, but despite a higher total number of species identified, there was little overlap and the two approaches likely targeted different species (Table 1 and Table 3). Both methods found similar numbers overall, though, so there appears to be a tradeoff between reducing labor (eDNA) and reducing cost/resources (traditional sampling), and ultimately, both approaches may be needed. For examples, species caught in nets are often large temporary visitors to flowers reducing the likelihood of leaving eDNA. Species found in eDNA sampling but not traditional trapping may represent nocturnal species or species too small to catch with traditional methods. Many of the arthropods found through eDNA Nanopore metabarcoding represented families that live on or around the flower head of a plant, contributing higher amounts of eDNA or DNA directly to the flower surface than other pollinators might. This is in line with other eDNA studies that missed large pollinators and found greater abundance of non-hymenopteran species from flower eDNA, potentially due to the difficulty in being able to remove eggs and other microscopic larvae from flowers before DNA extraction (Thomsen & Sigsgaard 2019; Johnson *et al.* 2023; Newton *et al.* 2023). eDNA sampling may be biased towards species that live on or around the flower as opposed to visiting pollinators. When looking at all high quality sequencing, the two survey methods shared 5.8 % of the same species, and this is in line with previous studies that found 5.3 % overlap between eDNA species and visual surveys (Newton *et al.* 2023) and 5 % overlap between eDNA species and camera traps (Johnson *et al.* 2023). The concordance with other studies supports our findings that multiplexed eDNA Nanopore metabarcoding is a valid approach for eDNA studies, however, the non-overlap between traditional and eDNA methods suggests it is important to include multiple sampling methods with eDNA analysis like physical or camera trapping to capture the full list of insects that use a specific flowering plant.

### Insights for pollinator taxa and native/non-native plants

The Empire State Native Pollinator Survey (ESNPS) was a four-year effort by scientists across the state to determine the current distribution and conservation status of selected pollinators. They focused on a core list of focal taxa, and we used this list along with the results from the traditional sampling to gain insight into New York insect pollinators (Supplemental Table S3). Traditional sampling found species from three focal taxa families listed in the Native Pollinator survey (Supplemental table S3), including the hover flies, *Syrphidae,* the solitary ground-nesting bees*, Andrenidae,* and the social bees, *Apidae* (Table 1). eDNA also identified multiples species found in the ENSPS, the hover flies, *Syrphidae* (also found in traditional sampling), the flower moths, *Noctuidae,* and some species of flower longhorn beetles, *Cerambycidae*, though the specific species listed in the ENSPS of flower longhorn beetle was not found. Between both methods, we found five families listed in the survey. In addition to identifying important pollinators from the ESNPS, we identified other arthropods from eDNA that may also be contributing to pollination. For example, the gall midge, *Ozirhincus millefolii*, is a plant parasite, but may also contribute to pollination through the transfer of pollen between flowers (Dorchin *et al.* 2015).

We also wanted to look at arthropod – plant network relationships in terms of native versus non-native flowers across the meadows. The majority of insect species found from eDNA were found only in native plants and not in exotic species (60 %). We found only eight (22.8 %) instances of an insect occurring solely on a non-native flower. There is increasing evidence of declining native plant species and shifts in their distribution (Mola *et al.* 2021), which has a large impact on native arthropod species that have evolved specialized and narrow plant relationships (Mola *et al.* 2021; Newton *et al.* 2023). From the eDNA data this may also be the case, where all arthropod species found that were part of the focal taxa from the ESNPS were found only on native plant species (Table 3). The hover fly, *Toxomerus geminatus,* was recovered from native bergamot (*Monarda fistulosa).* The flower moth, *Schinia arcigera*, was found on native flat-topped golden rod (*Euthamia graminifolia*). Finally, the flower longhorn beetles, *Typocerus velutinus* and *Tetraopes quinquemaculatus* were found on flat-top white aster (*Aster umbellatus*) and milkweed (*Asclepias tuberosa*), respectively. These results show that eDNA sampling of flowers can uncover arthropod-plant relationships that may be missed through traditional sampling and underscores the importance of maintaining native plant diversity to increase native pollinator diversity, especially in the face of climate change.

### Conclusion

The results we present show the validity of multiplexed eDNA Nanopore sequencing to identify animal species from flower heads. Nanopore sequencing using the MinION is fast and portable. By incorporating unique tags into the initial primer design and using the reduced pore flongle flow cells, allowing for dozens of samples to be multiplexed in a single run, nanopore sequencing can also be extremely affordable. Our study is the first to show the feasibility of multiplexed Nanopore sequencing from flower eDNA and our methodology can be utilized in labs or in the field. The method is efficient and non-invasive compared to traditional techniques and analysis can be performed months after specimen collection. Freezing is the only requirement for preservation, and flower heads are small, lightweight and can be easily transported –meaning insect biodiversity can be studied with relatively low impact. However, our study supports findings from others that eDNA sequencing alone is not enough to identify all pollinators visiting flowers. Combining multiplexed eDNA Nanopore metabarcoding with another non-invasive method like camera traps (Johnson *et al.* 2023) will likely maximize species recovery and identification. Our study found eDNA from a large variety animals on flowers, providing many different species than identified from traditional sampling. Species included pollinators, predators, and phytophagous species while being able to link each with a specific flower species, location, and sampling time. This allowed us to show that eDNA metabarcoding of wildflowers can successfully be applied to conservation and biodiversity tracking. We show that large floral-diverse meadows like BM hold high insect species richness and increasing native flowers is important attracting native pollinators. Future research may sample the full catalogue of native flower species in NY to inform which plants are most important for recruiting and supporting important native pollinators. Additionally, this approach can provide diagnostic possibilities for protected lands by providing early warnings for the presence of invasive species. Multiplexed eDNA nanopore metabarcoding is a reliable and cost-effective method for uncovering arthropod diversity, and in combination with other visual methods can provide insight into arthropod-plant networks and how they respond to an increasingly changing climate.

## Acknowledgements

We thank the Mianus River Gorge for access to study sites. We thank the Pollinator Network at Cornell for help in identifying collected insects. We thank Leah Cass, Emily Bosch Mak Boylan, Lucas Andujar, Serena Feldman, and Jean-Luc Plante for their assistance in the field. This work was funded by an Undergraduate Research Grant from Purchase College, SUNY, a 2021-22 Craddock Scholar award from SUNY Purchase, and the Rusticus Garden Club.

## Author Contributions

Conceived and designed the experiments: SEH BV CN. Field work and sample collection: SEH AW CN BV SL MB. Performed the experiments: AW SEH. Analyzed the data: AW SEH MB SL. Contributed reagents/materials/analysis tools: SEH CN BV. Wrote original manuscript: SEH AW. Review and editing SEH AW CN. Visualization SEH AW CN. Resources SEH CN BV. Supervision SEH CN BV. Administration SEH CN. Funding SEH CN.

## Data Accessibility

All demultiplexed Nanopore sequencing data in fastq format (q ≥ 10) and full Blastn results have been uploaded and made available from the Dryad Digital Repository: DRYAD entry doi:xx.xxx/dryad.XXXX.

**-** Supplemental Table S1: Complete Plant Collection Data
**-** Supplemental Table S2: Min 100 length and 98% ID Blast results
**-** Supplemental Table S3: ESNPS focal taxa

## Notes

### Competing Interest Statement

The authors have declared no competing interest.

