## Supplemental Table S1 for "Efficient Multiplexing of Pollinator Metabarcodes Using Oxford Nanopore MinION Sequencing: Insights for Meadow Management from Floral Environmental DNA"

Supplemental Table S1: List and frequency of all flowers collected for this study.

A. Wildflowers Collected June 28, 2021 at Mianus River Gorge Preserve

| Lockwood North | Frequency | Common Name | Scientific | Family | NY Native Status |
| --- | --- | --- | --- | --- | --- |
|  | 1 | whorled coreopsis | *Coreopsis verticillata* | *Asteraceae* | Native |
|  | 3 | milkweed | *Asclepias tuberosa* | *Apocynaceae* | Native |
|  | 1 | narrow leaf mint | *Pycnanthemum tenuifolium* | *Lamiaceae* | Native |
|  | 1 | field bindweed | *Convolvulaceae sp.* | *Convolvulaceae* | Exotic |
|  | 1 | large hop clover | *Trifolium aureum* | *Fabaceae* | Exotic |
|  | 2 | common yarrow | *Achillea millefolium* | *Asteraceae* | Native |
|  | 1 | red clover | *Trifolium pratense* | *Fabaceae* | Exotic |
|  | 1 | mullein | *Verbascum spp.* | *Scrophulariaceae* | Native |
|  | 1 | scarlet bergamot | *Monarda didyma* | *Lamiaceae* | Native |
|  | 12 | Number of Specimens |  |  |  |
|  | 9 | Number of Species |  |  |  |
| Big Meadow | Frequency | Common Name | Scientific | Family |  |
|  | 2 | blackeyed susan | *Rudbeckia hirta* | *Asteraceae* | Exotic |
|  | 3 | milkweed | *Asclepias tuberosa* | *Apocynaceae* | Native |
|  | 1 | common st. john’s wort | *Hypericum perforatum* | *Hypericaceae* | Exotic |
|  | 5 | dogbane (indian hemp) | *Apocynum cannabinum* | *Apocynaceae* | Native |
|  | 3 | large hop clover | *Trifolium aureum* | *Fabaceae* | Exotic |
|  | 6 | red clover | *Trifolium pratense* | *Fabaceae* | Exotic |
|  | 3 | tufted vetch | *Achillea millefolium* | *Asteraceae* | Exotic |
|  | 2 | upright yellow wood sorrel | *Oxalis stricta* | *Oxalidaceae* | Native |
|  | 3 | white clover | *Trifolium repens* | *Fabaceae* | Exotic |
|  | 1 | common yarrow | *Achillea millefolium* | *Asteraceae* | Native |
|  | 30 | Number of specimens collected |  |  |  |
|  | 10 | Number of Species |  |  |  |
|  | 1 | Number of Unknown |  |  |  |
| Captain’s Woods | Frequency | Common Name | Scientific | Family |  |
|  | 4 | daisy fleabane | *Erigeron spp* | *Asteraceae* | Native |
|  | 1 | yarrow | *Achillea millefolium* | *Asteraceae* | Native |
|  | 1 | Carolina horse nettle | *Solanum carolinense* | *Solanaceae* | Native |
|  | 2 | milkweed | *Asclepias spp.* | *Apocynaceae* | Native |
|  | 2 | white avens | *Geum canadense* | *Rosaceae* | Native |
|  | 10 | Number of Specimens Collected |  |  |  |
|  | 5 | Number of Species Present |  |  |  |

B. Wildflowers collected August 11, 2021 at Mianus River Gorge Preserve

| **Lockwood North** | Frequency | Common Name | Scientific | Family | New York Native Status |
| --- | --- | --- | --- | --- | --- |
|  | 2 | American ground-nut, common | *Apios americana* | *Fabaceae* | Native |
|  | 1 | dodder, field | *Cuscuta campestris* | *Convolvulaceae* | Native |
|  | 1 | goldenrod | *Goldenrod spp.* | *Asteraceae* | Native |
|  | 1 | hedge bindweed, false | *Calystegia sepium* | *Convoluceae* | Exotic |
|  | 1 | large hop clover | *Trifolium campestre* | *Fabaceae* | Exotic |
|  | 3 | moth muellin | *Verbascum blattaria* | *Scrophulariaceae* | Exotic |
|  | 1 | New York ironweed | *Vernonia noveboracensis* | *Asteraceae* | Native |
|  | 1 | scarlet bee-balm | *Monarda didyma* | *Lamiaceae* | Native |
|  | 1 | naked tick-trefoil | *Hylodesmum nudiflorum* | *Fabaceae* | Native |
|  | 1 | upright yellow wood sorrel | *Oxalis stricta* | *Oxalidaceae* | Native |
|  | 2 | Virginia mountain-mint | *Pycnanthemum virginianum* | *Lamiaceae* | Native |
|  | 2 | wild basil | *Clinopodium vulgare* | *Lamiaceae* | Exotic |
|  | 1 | wild bergamont | *Monarda fistulosa* | *Lamiaceae* | Native |
|  | 6 | wild carrot | *Daucus carota* | *Apiaceae* | Exotic |
|  | 1 | woodland sunflower | *Helianthus divaricatus* | *Asteraceae* | Native |
|  | 2 | yarrow | *Achillea millefolium* | *Asteraceae* | Native |
|  | 27 | Number of Specimens |  |  |  |
|  | 1 | Number of Unknown |  |  |  |
|  | 1 | Number MIA |  |  |  |
|  | 17 | Number of Species |  |  |  |
| **Big Meadow** | Frequency | Common Name | Scientific Name | Family |  |
|  | 2 | black-eyed susan | *Rudbeckia hirta* | *Asteraceae* | Exotic |
|  | 2 | milkweed | *Asclepias tuberosa* | *Apocynaceae* | Native |
|  | 1 | common self-heal | *Prunella vulgaris* | *Lamiaceae* | Exotic |
|  | 1 | common toad flax | *Linaria vulgaris* | *Plantaginaceae* | Exotic |
|  | 1 | flat-top golden-seal | *Hydrastis canadensis* | *Ranunculaceae* | Native |
|  | 2 | flat-top white aster | *Aster umbellatus* | *Asteraceae* | Native |
|  | 1 | flat-topped goldenrod | *Euthamia graminifolia* | *Asteraceae* | Native |
|  | 1 | hedge bindweed, false | *Calystegia sepium* | *Convoluceae* | Exotic |
|  | 2 | large hop clover | *Trifolium campestre* | *Fabaceae* | Exotic |
|  | 4 | red clover | *Trifolium pratense* | *Fabaceae* | Exotic |
|  | 1 | red thistle ( swamp thistle) | *Cirsium muticum* | *Asteraceae* | Native |
|  | 1 | soapwort | *Saponaria spp.* | *Caryophyllaceae* | Native |
|  | 4 | tufted vetch | *Vicia cracca* | *Fabaceae* | Exotic |
|  | 1 | unknown sunflower | *Helianthus spp.* | *Asteraceae* | N/A |
|  | 3 | upright yellow wood sorrel | *Oxalis stricta* | *Oxalidaceae* | Native |
|  | 2 | wild basil | *Clinopodium vulgare* | *Lamiaceae* | Exotic |
|  | 9 | wild bergamont | *Monarda fistulosa* | *Lamiaceae* | Native |
|  | 5 | wild carrot (queen anne's lace) | *Daucus carota* | *Apiaceae* | Exotic |
|  | 5 | yarrow, common | *Achillea millefolium* | *Asteraceae* | Native |
|  | 48 | Number of Specimens |  |  |  |
|  | 1 | Number of MIA |  |  |  |
|  | 19 | Number of Species |  |  |  |
| **Captain’s Woods** | Frequency | Common Name | Scientific | Family |  |
|  | 2 | American hog-peanut | *Amphicarpaea bracteata* | *Fabaceae* | Native |
|  | 1 | annual fleabane | *Errigeron annus* | *Asteraceae* | Native |
|  | 2 | Carolina horsenettle | *Solanum carolinense* | *Solanceae* | Native |
|  | 2 | field dodder | *Cuscuta campestris* | *Convolvulaceae* | Native |
|  | 8 | flat-topped goldenrod | *Euthamia graminifolia* | *Asteraceae* | Native |
|  | 1 | flat-top white aster | *Aster umbellatus* | *Asteraceae* | Native |
|  | 3 | common wrinkle-leaved goldenrod | *Solidago rugosa* | *Asteraceae* | Native |
|  | 1 | horse weed | *Erigeron canadensis* | *Asteraceae* | Native |
|  | 1 | jewel weed | *Impatiens capensis* | *Balsaminaceae* | Native |
|  | 3 | joe pie weed | *Eutrochium maculatum* | *Asteraceae* | Native |
|  | 1 | lesser burdock | *Arctium minus* | *Asteraceae* | Exotic |
|  | 7 | New York ironweed | *Vernonia noveboracensis* | *Asteraceae* | Native |
|  | 1 | ox -eye daisy | *Leucanthemum vulgare* | *Asteraceae* | Exotic |
|  | 1 | pokeweed, American | *Phytolacca americana* | *Phytolaccaceae* | Native |
|  | 1 | swamp sunflower | *Helianthus angustifolius* | *Asteraceae* | Native |
|  | 1 | Virginia mountain-mint | *Pycnanthemum virginianum* | *Lamiaceae* | Native |
|  | 1 | yarrow, common | *Achillea millefolium* | *Asteraceae* | Native |
|  | 37 | Number of Specimens Collected |  |  |  |

C. Wildflowers collected September 7,2021 at Mianus River Gorge

| **Captain** **Woods** | Frequency | Common Name | Scientific Name | Family | New York Native Status |
| --- | --- | --- | --- | --- | --- |
|  | 4 | American aster | *Aster spp.* | *Asteraceae* | Native |
|  | 2 | arrow leaved tear thumb | *Persicaria sagittata* | *Polygonaceae* | Exotic |
|  | 6 | calico aster | *Aster hirsuticaulis* | *Asteraceae* | Native |
|  | 1 | climbing house buckwheat | *Fagopyrum spp.* | *Polygonaceae* | Exotic |
|  | 5 | common wrinkle-leaved goldenrod | *Solidago rugosa* | *Asteraceae* | Native |
|  | 1 | dayflower | *Commelina communis* | *Commelinaceae* | Exotic |
|  | 2 | field dodder | *Cuscuta campestris* | *Convolvulaceae* | Native |
|  | 7 | flat-topped goldenrod | *Euthamia graminifolia* | *Asteraceae* | Native |
|  | 2 | flat-top white aster | *Aster spp.* | *Asteraceae* | Native |
|  | 1 | goldenrod spp | *Solidago sp* | *Asteraceae* | Native |
|  | 1 | jewel weed | *Impatiens capensis* | *Balsaminaceae* | Native |
|  | 3 | New York ironweed | *Vernonia noveboracensis* | *Asteraceae* | Native |
|  | 1 | pokeweed, American | *Phytolacca americana* | *Phytolaccaceae* | Native |
|  | 2 | sunflower | *Helianthus spp* | *Asteraceae* | N/A |
|  | 38 | Number of Spcecimens Collected |  |  |  |
|  | 2 | Number of MIA |  |  |  |
|  | 35 | Number of Spcecimens Collected within Grid |  |  |  |
|  | 3 | Number of Specimens Collected outside of Grid |  |  |  |
|  | 13 | Number of Species |  |  |  |
| **Big** **Meadow** | Frequency | Common Name | Scientific | Family |  |
|  | 1 | American aster | *Aster spp.* | *Asteraceae* | Native |
|  | 3 | calico aster | *Aster hirsuticaulis* | *Asteraceae* | Native |
|  | 1 | clover | *Trifolium spp* | *Fabaceae* | Exotic |
|  | 1 | common toad flax | *Linaria vulgaris* | *Plantaginaceae* | Native |
|  | 9 | common wrinkled-leaf goldenrod | *Solidago rugosa* | *Asteraceae* | Native |
|  | 3 | common yarrow | *Achillea millefolium* | *Asteraceae* | Native |
|  | 3 | flat-top goldenrod | *Euthamia graminifolia* | *Asteraceae* | Native |
|  | 6 | flat-top white aster | *Aster umbellatus* | *Asteraceae* | Native |
|  | 1 | flax-leaved aster | *Aster linariifolius* | *Asteraceae* | Native |
|  | 6 | late purple aster | *Aster patens* | *Asteraceae* | Native |
|  | 2 | red (scarlet) clover | *Trifolium pratense* | *Fabaceae* | Exotic |
|  | 1 | silverrod (aka white goldenrod) | *Solidago bicolor* | *Asteraceae* | Native |
|  | 1 | tuffted vetch | *Achillea millefolium* | *Asteraceae* | Exotic |
|  | 1 | Virginia mountain mint | *Pycnanthemum virginianum* | *Lamiaceae* | Native |
|  | 4 | wild basil | *Clinopodium vulgare* | *Lamiaceae* | Exotic |
|  | 2 | wild bergamont | *Monarda fistulosa* | *Lamiaceae* | Native |
|  | 3 | wild carrot | *Daucus carota* | *Apiaceae* | Exotic |
|  | 48 | Number of Specimens Collected |  |  |  |
|  | 45 | Number of Specimens from Grid |  |  |  |
|  | 3 | Number of Specimens Present outside of Grid |  |  |  |
|  | 17 | Number of Species |  |  |  |
| **Lockwood** **North** | Frequency | Common Name | Scientific | Family |  |
|  | 2 | American hog Peanut | *Amphicarpaea bracteata* | *Fabaceae* | Native |
|  | 1 | arrow-leaved tearthumb | *Persicaria sagittata* | *Polygonaceae* | Native |
|  | 2 | calico aster | *Aster hirsuticaulis* | *Asteraceae* | Native |
|  | 3 | common wrinkled-leaf goldenrod | *Solidago rugosa* | *Asteraceae* | Native |
|  | 1 | dark mullein | *Verbascum spp.* | *Scrophulariaceae* | Exotic |
|  | 5 | flat-topped goldenrod | *Euthamia graminifolia* | *Asteraceae* | Native |
|  | 3 | late purple aster | *Symphyotrichum patens* | *Asteraceae* | Native |
|  | 1 | narrow leaf mountain mint | *Pycnanthemum tenuifolium* | *Lamiaceae* | Native |
|  | 1 | scarlett bee balm/ bergamont | *Monarda didyma* | *Lamiaceae* | Native |
|  | 1 | sunflower sp | *Helianthus. spp* | *Asteraceae* | Native |
|  | 1 | oriental lady's thumb smartweed | *Persicaria longiseta* | *Polygonaceae* | Exotic |
|  | 1 | upright yellow wood sorrel | *Oxalis stricta* | *Oxalidaceae* | Native |
|  | 1 | white wood-aster | *Aster divaricatus* | *Asteraceae* | Native |
|  | 1 | whorled coreopsis | *Coreopsis verticillata* | *Asteraceae* | Native |
|  | 2 | wild basil | *Clinopodium vulgare* | *Lamiaceae* | Exotic |
|  | 5 | wild carrot | *Daucus carota* | *Apiaceae* | Exotic |
|  | 31 | Number of Specimens Collected |  |  |  |
|  | 16 | Number of Species |  |  |  |
