## Supplemental Table S3 for "Efficient Multiplexing of Pollinator Metabarcodes Using Oxford Nanopore MinION Sequencing: Insights for Meadow Management from Floral Environmental DNA"

Supplemental Table S3: List of focal taxa from the Empire State Native Pollinator Survey. Focal taxa determined by the Empire State Native Pollinator Survey from 2016 – present. Tablre reproduced from https://www.nynhp.org/projects/pollinators/.

| **Order and CommonNames** | **Scientific Names** |
| --- | --- |
| **Hymenoptera** |  |
| Bumble bees and long-horned bees | Apidae: *Bombus, Melissodes* |
| Mining bees | Andrenidae: *Andrena, Calliopsis* |
| Leafcutter bees | Megachilidae: *Megachile, Osmia* |
| Oil bees | *Macropis, Melitta* |
| A cuckoo bee | *Epeoloides pilosula* |
| **Diptera** |  |
| Bee flies | Bombyliidae: *Bombylius* |
| Saproxylic (decaying wood) hover flies | Syrphidae: ~80 species in two subfamilies |
| **Coleoptera** |  |
| Flower longhorn beetles | Cerambycidae: *Lepturinae* |
| Hairy flower scarabs | Scarabeidae: *Trichiotinus* |
| **Lepidoptera** |  |
| Hawk (sphinx) moths | Sphingidae: 26 species that feed as adults |
| Flower moths | Noctuidae: *Schinia* |
